# Benchmarking the AlphaGenome Variant Impact score against SpliceAI on post-release ClinVar classifications and experimental splicing assays

**DOI:** 10.64898/2026.10.11.758362

**Authors:** Ozge A. Cavus, Aysegul Kuskucu

## Abstract

The AlphaGenome Variant Impact (AVI) score ranks variants by predicted impact, and its developers report large gains over other genome-wide scores for intronic ClinVar variants. We tested AVI on two kinds of ground truth the developers did not use, under an analysis plan set before scoring: ClinVar classifications made after the developers’ ClinVar release (87,666 rare single-nucleotide variants outside AVI’s training chromosomes), and six splicing assays whose variants no predictor chose. In intronic variants at least 3 bases from an exon (449 pathogenic or likely pathogenic, 28,188 benign or likely benign), AVI reached an area under the receiver operating characteristic curve (AUROC) of 0.957. It did not differ detectably from SpliceAI (difference −0.005, 95% interval −0.017 to 0.006) or Pangolin (−0.006, −0.018 to 0.007); both intervals exclude differences larger than about 0.02. AVI had a higher AUROC than CADD (+0.071). In deep intronic variants it trailed SpliceAI (−0.070, −0.118 to −0.026). Under an adapted ClinGen calibration, AVI gave strong evidence to 52% of pathogenic variants on the pooled test set, but its levels did not hold in every region. It gave benign evidence to 24% of benign variants, against 90% for SpliceAI. On the assays, AVI ranked splice-altering variants below raw SpliceAI (pooled difference −0.089, −0.129 to −0.049) and did not differ from masked SpliceAI (+0.003). SpliceAI2, released during the study, led AVI on both kinds of ground truth. AVI did not outperform splicing predictors on AUROC and gave little benign evidence. AlphaGenome outputs are provided for research use, not for diagnostic decisions.

## Introduction

Most variants found in rare-disease sequencing lie outside protein-coding sequence or do not change an amino acid. Their interpretation leans heavily on computational predictions. Under the ACMG/AMP framework [Richards et al. 2015], predictions enter as supporting evidence for pathogenicity (PP3) or benignity (BP4). The ClinGen Sequence Variant Interpretation working group showed how a predictor can be calibrated so that a score interval carries supporting, moderate or strong evidence, given a prior probability of pathogenicity [Pejaver et al. 2022]. Calibrated thresholds have since been published for SpliceAI [Walker et al. 2023], whose scores are now used routinely for variants that may affect splicing.

AlphaGenome predicts thousands of molecular read-outs from DNA sequence, including splice-site use, splice junctions, RNA expression and chromatin accessibility [Avsec et al. 2026]. The AlphaGenome Variant Impact (AVI) score combines in silico mutagenesis predictions from AlphaGenome with AlphaMissense [Cheng et al. 2023], two conservation scores and protein-coding annotations in one supervised model [Cheng et al. 2026]. AVI was trained to separate variants with a gnomAD v4.1 filtering allele frequency below 0.1% from those above it, a proxy for selection rather than a clinical label. Its developers evaluated it on ClinVar classifications from the release of 2025-06-15. For intronic single-nucleotide variants they report an area under the precision-recall curve (AUPRC) of 0.76, against 0.44 for the next-best genome-wide model [Cheng et al. 2026].

Two questions remain open. First, an evaluation on ClinVar labels that were available during development, and on benchmarks used to choose between model versions, can be optimistic. Second, ClinVar labels for non-coding variants are partly shaped by predictions: a variant flagged by SpliceAI can be more likely to be tested and classified. Labels of this kind can favour the predictors that helped produce them, and they can make any newer model look weaker or stronger than it is. Neither question can be settled by the developers’ own benchmarks.

We therefore evaluated AVI on two kinds of ground truth that the developers did not use. The first is ClinVar classifications made after the release the developers used. The second is splicing assays that measured every possible variant, or a random set of variants, in an exon and its flanks; one library also measured variants observed in a population database. In none of these assays did a predictor choose which variants were tested. We compared AVI with splicing predictors (SpliceAI, Pangolin), a genome-wide score (CADD v1.7), conservation scores and a baseline based on distance to the exon, and we measured how AVI scores behave under the ClinGen calibration framework. The analysis plan, variant sets and metrics were written and committed to the project repository before any score was retrieved.

## Results

### Study sets

Part 1 started from the 1,762,164 variants with a pathogenic or likely pathogenic (P/LP) or benign or likely benign (B/LB) classification in the ClinVar release of October 2026 (Fig. 1; Supplementary Table 1). We removed variants already classified P/LP or B/LB at or before the 2025-06-15 release (1,426,176), variants in the genes or at the positions of the developers’ benchmark datasets (Methods) or submitted by an organisation that supplied the developers’ cases (123,767), 818 variants at the position of an older record of unresolved identity, and 17 identity collisions. This left 211,386 new labels (49,280 P/LP, 162,106 B/LB). The main analysis set kept single-nucleotide variants in intronic, untranslated or synonymous positions, with at least one review star, with an HGVS description on the gene’s MANE Select transcript, on the developers’ validation chromosomes and on those they used neither for training nor for validation (‘untouched’; Methods), and rare in gnomAD v4.1: 87,666 variants (4,453 P/LP, 83,213 B/LB). Variants on training chromosomes (40,403) and common variants (4,111) were kept for supporting analyses.

**Figure 1.**
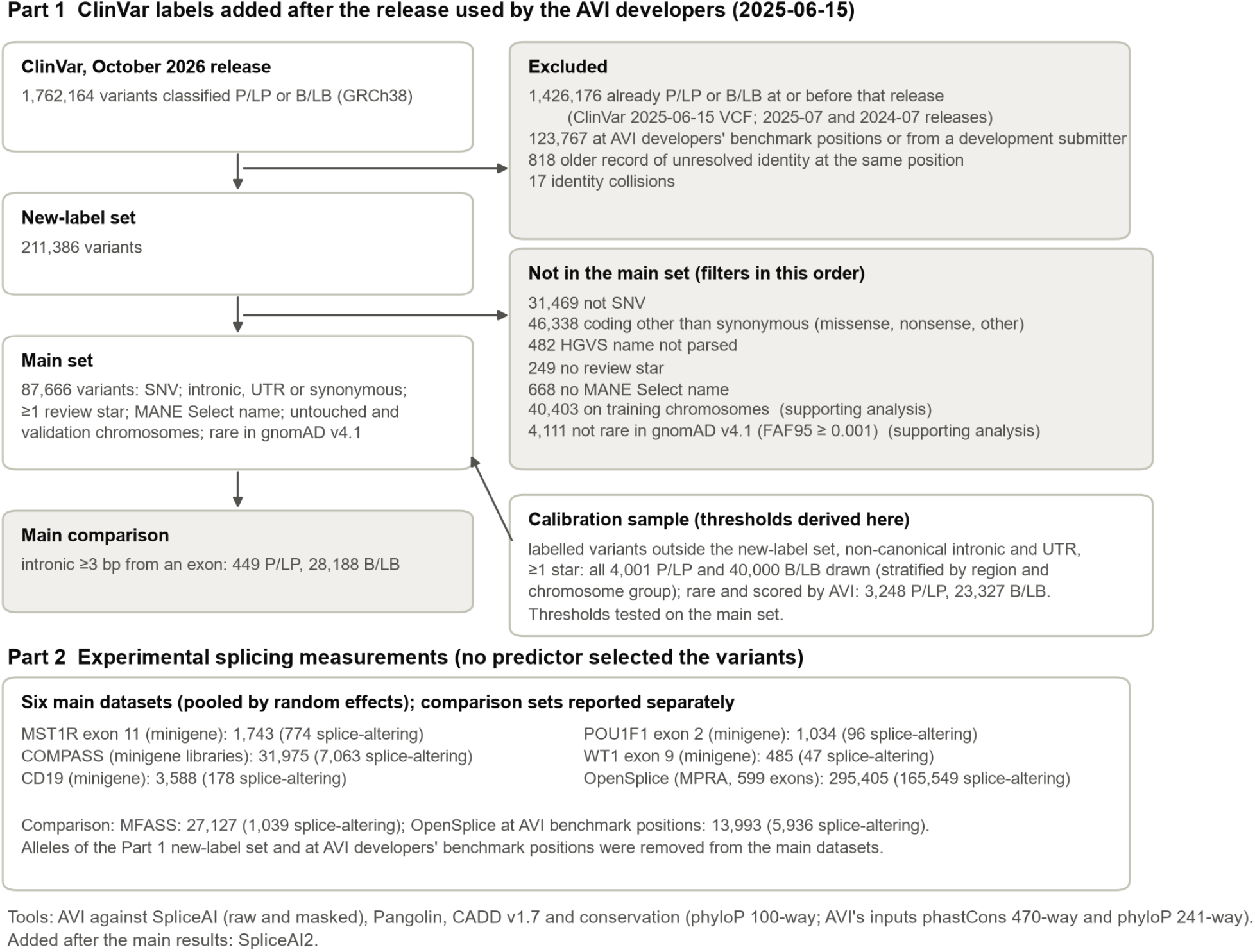
Study design. Part 1 uses ClinVar classifications first made after the ClinVar release used by the AVI developers (2025-06-15; AVI itself was trained on population frequency, not on ClinVar); thresholds for clinical evidence are derived on a separate calibration sample of labelled variants outside the new-label set and tested on the main set. Part 2 uses experimental splicing measurements in which no predictor selected the variants. Counts are variants (Part 1), with the filters from the new-label set to the main set applied in the order listed, or SNV assay variants with the number measured as splice-altering (Part 2).

The main comparison region was intronic variants at least 3 bases from the exon, which pools the donor region (+3 to +6), the acceptor region (−3 to −20), near intronic variants (+7 to +50 and −21 to −50) and deep intronic variants (beyond 50 bases): 449 P/LP in 266 genes and 28,188 B/LB. AVI, CADD and SpliceAI2 scored every main-set variant; precomputed SpliceAI scored 99% (Supplementary Table 2).

Part 2 used six main datasets (334,230 variants in total; Supplementary Table 3): saturation minigene assays of POU1F1 exon 2 (1,034 variants, 96 splice-altering) [Gergics et al. 2021], WT1 exon 9 (485; 47) [Smith et al. 2023], MST1R exon 11 (1,743; 774) [Braun et al. 2018] and CD19 exons 1–3 (3,588; 178) [Cortés-López et al. 2022], and two multiplexed libraries, COMPASS (31,975; 7,063) [Koplik et al. 2025], which holds random variants and an ExAC subset (Methods), and OpenSplice (295,405; 165,549) [Quarantani et al. 2026]. MFASS (27,127; 1,039) [Chong et al. 2019] was analysed as a comparison dataset because the developers of AlphaGenome and of Pangolin report results on it.

### Discrimination on new clinical labels

In the main region AVI reached an area under the receiver operating characteristic curve (AUROC) of 0.957 (0.944–0.969), SpliceAI 0.962 (0.949–0.973), Pangolin 0.962 (0.947–0.975) and CADD 0.886 (0.864–0.907) (Fig. 2a). On the variants both tools scored, AVI did not differ detectably from SpliceAI (paired difference −0.005, −0.017 to 0.006) or Pangolin (−0.006, −0.018 to 0.007); the intervals exclude differences larger than about 0.02. AVI had a higher AUROC than CADD (+0.071, 0.053 to 0.092). A baseline that ranks variants by distance to the nearest exon boundary reached 0.663 (0.609–0.714), below every other score.

**Figure 2.**
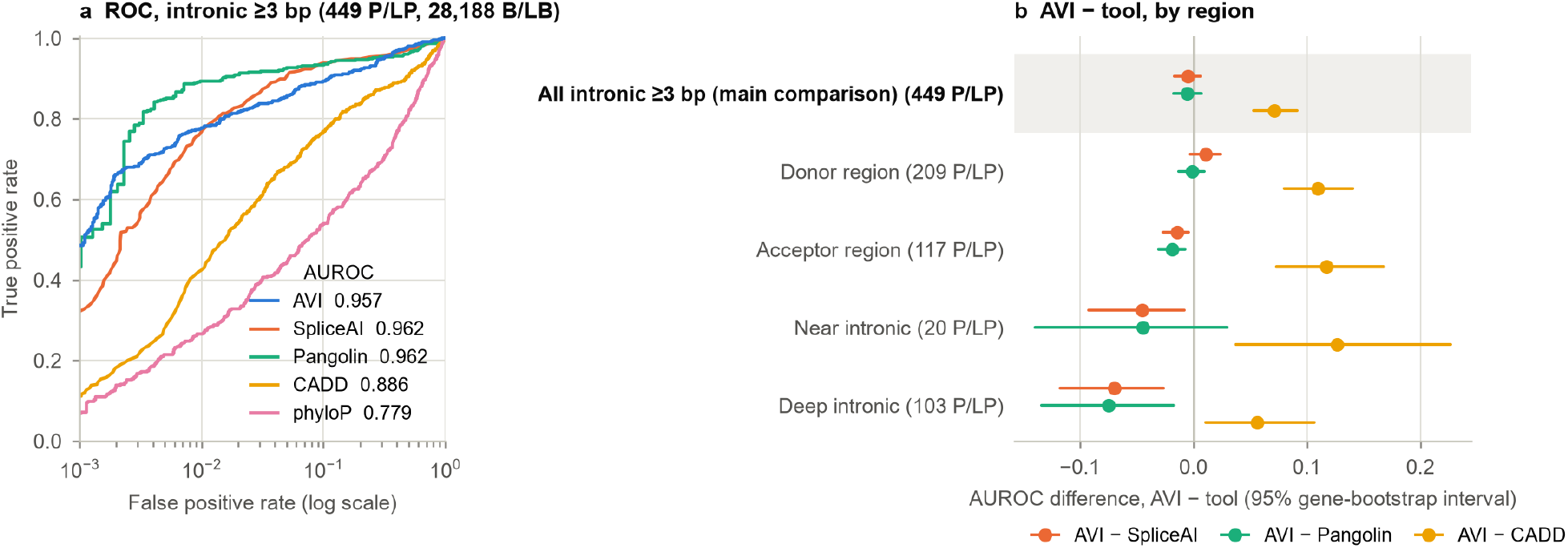
Discrimination on ClinVar labels added after the release used by the AVI developers (Part 1). (a) ROC curves for intronic variants at least 3 bases from an exon in the main set (449 P/LP, 28,188 B/LB); false positive rate on a log scale to show the high-specificity range. Pangolin was run on all P/LP and a region-stratified sample of 4,000 B/LB; its curve and AUROC use inverse-sampling weights, and its curve is stepped at low false positive rates because fewer B/LB were scored. phyloP, 100-way vertebrate phyloP. (b) Paired AUROC differences, AVI minus each tool, on the variants both tools score; the main comparison (all intronic variants ≥3 bp) is shown first. Intervals are 95% gene-cluster bootstrap intervals (2,000 replicates).

By region (Fig. 2b; Supplementary Table 4), AVI did not differ from SpliceAI in the donor region (+0.011, −0.003 to 0.024) and was slightly behind it in the acceptor region (−0.014, −0.027 to −0.004) and in near intronic variants (−0.045, −0.092 to −0.008; 20 P/LP). The largest difference was in deep intronic variants, more than 50 bases from the exon (103 P/LP, 1,572 B/LB). There AVI reached 0.820 (0.759– 0.872) and trailed SpliceAI (−0.070, −0.118 to −0.026), Pangolin (−0.075, −0.134 to −0.017) and SpliceAI run locally with a 4,999-base window (−0.075, −0.128 to −0.024). Against CADD, AVI was ahead in the donor (+0.110), acceptor (+0.117), near intronic (+0.127) and deep intronic (+0.056, 0.011 to 0.107) regions and in synonymous variants (+0.043, 0.015 to 0.077). In canonical splice sites (25 B/LB) and untranslated regions (12 P/LP) the intervals were too wide to tell the tools apart. AVI outperformed its own conservation inputs by 0.186 (phastCons 470-way; 0.161 to 0.211) and 0.187 (Cactus 241-way phyloP; 0.161 to 0.210) in the main region.

The main-region results held in every supporting analysis (Supplementary Table 5). Across all frequencies, variants present in gnomAD, training chromosomes, variants with at least two review stars, the stricter exclusion of older unresolved records, the calibration sample, and the subsets in which a ClinVar submission did or did not mention a splicing predictor, AVI minus SpliceAI ranged from −0.017 to +0.005 and AVI minus CADD from +0.069 to +0.095. AVI was not better on the chromosomes it was trained on (AVI minus SpliceAI +0.005, −0.017 to 0.026). Under the earlier, stricter exclusion of older records (Methods), which kept 235 P/LP and 18,505 B/LB in the main region, AVI minus SpliceAI on the variants both scored was −0.004 (−0.019 to 0.011).

Labels could favour SpliceAI if the submitters used it. Among variants whose ClinVar submissions did not mention a splicing predictor, AVI and SpliceAI did not differ in the main region (0.000, −0.013 to 0.013); where a submission did mention one, the difference was −0.017 (−0.040 to 0.008; 131 P/LP, 54 B/LB), the widest of the supporting analyses. In deep intronic variants without a predictor mention (71 P/LP), the point estimates were smaller than in the full set and the intervals wider: −0.040 (−0.096 to 0.011) against SpliceAI and −0.041 (−0.106 to 0.024) against Pangolin. Where a submission did mention a predictor (32 deep intronic P/LP, 16 B/LB), the difference to SpliceAI was −0.108 (−0.230 to 0.011). The deep intronic difference also appeared in older labels: in the calibration sample (506 P/LP) AVI minus SpliceAI was −0.043 (−0.086 to −0.001), and on the training chromosomes −0.032 (−0.112 to 0.047).

### Calibration under the ClinGen framework

Thresholds for each evidence level were derived on a separate calibration sample of labelled variants outside the new-label set (for AVI, 3,248 P/LP and 23,327 B/LB) and tested on the main set’s non-canonical intronic and untranslated variants, the pooled test set (461 P/LP and 28,858 B/LB for AVI; Fig. 3; thresholds in Supplementary Table 6, likelihood ratios in Supplementary Table 7 and, in cumulative form, Supplementary Table 8). We call a level supported when the 95% interval of its likelihood ratio lies above the ClinGen target (below its reciprocal for benign levels).

**Figure 3.**
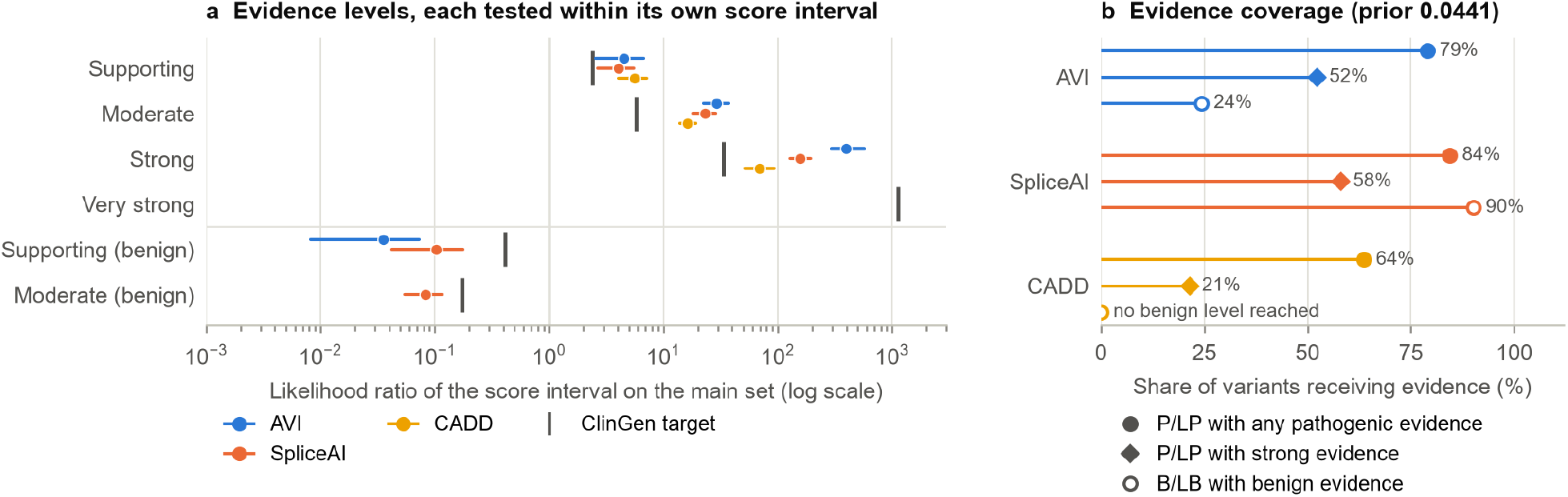
Calibration under the ClinGen framework (Part 1). (a) Interval likelihood ratios with 95% gene-bootstrap intervals for each evidence level, tested on the main set (non-canonical intronic and UTR variants) with thresholds derived on the calibration sample at a prior of 0.0441 (Methods). Vertical bars: ClinGen targets (2.406, 5.790, 33.53 and 1,124; benign levels the reciprocals). A level without a mark was not reached. (b) Share of P/LP receiving any pathogenic evidence and strong evidence, and of B/LB receiving any benign evidence. Likelihood ratios in (a) are not compared across tools, because each tool’s intervals differ; the shares in (b) are. Likelihood ratios of different tools are measured on different score intervals and are not comparable. The shares of variants receiving evidence are given in Table 1 (Fig. 3b).

**Table 1.** Share of variants receiving evidence on the pooled test set (prior 0.0441). SpliceAI2 was added as a comparator after the main results (Additional analyses).

| Tool | P/LP with any pathogenic evidence | P/LP with strong evidence | B/LB with any benign evidence |
| --- | --- | --- | --- |
| AVI | 79% | 52% | 24% |
| SpliceAI | 84% | 58% | 90% |
| SpliceAI2 | 86% | 67% | 91% |
| CADD | 64% | 21% | 0% |

For AVI, the supporting (likelihood ratio 4.51, 2.56–6.77), moderate (29.1, 22.5–37.3) and strong (397, 294–569) pathogenic levels and the supporting benign level (0.036, 0.008–0.074) were all supported. On this pooled test set the same held for SpliceAI, masked SpliceAI and CADD at every level each tool reached, and for SpliceAI2, a comparator added after the main results (Additional analyses). Masked SpliceAI sets to zero the predicted changes that strengthen annotated splice sites or weaken unannotated ones, as in the masked precomputed files. CADD reached no benign level. The very strong level (likelihood ratio 1,124) could not be demonstrated for any tool; it would require a far larger set of B/LB variants in the high-score window than this sample holds.

AVI therefore gives strong pathogenic evidence to slightly fewer P/LP variants than SpliceAI, and benign evidence to far fewer B/LB variants. The benign share depends on the prior. At a prior of 0.01 (Supplementary Table 9), AVI gave benign evidence to 0.85% of B/LB variants and SpliceAI to 83%; at 0.10, the shares were 51% and 90%.

Within regions the counts are smaller, and many levels could not be shown for any tool (Table 2; all tools in Supplementary Table 10). AVI’s moderate and strong pathogenic levels were supported in acceptor-region and near intronic variants (the latter from 20 P/LP), but none of its pathogenic levels was supported in donor-region or deep intronic variants. Its supporting benign level held in donor-region, acceptor-region and deep intronic variants but not in near intronic variants. In donor-region variants none of SpliceAI’s pathogenic levels was supported either, whereas in deep intronic variants its moderate and strong levels were. SpliceAI’s moderate benign level held in donor-region and acceptor-region variants; AVI reached a moderate benign level in no region. Untranslated regions held too few P/LP variants (12) to assess.

**Table 2.** Calibration by region on the main set: interval likelihood ratio (95% interval); S, supported; N, not supported. SpliceAI: precomputed raw scores, 50-base window. Each level is tested within its own score interval, so a higher level can be supported where a lower one is not. Untranslated regions (12 P/LP) are not shown.

| Region | Tool | P/LP<br>/<br>B/LB | Supporting<br>pathogenic | Moderate<br>pathogenic | Strong<br>pathogenic | Supporting<br>benign | Moderate<br>benign |
| --- | --- | --- | --- | --- | --- | --- | --- |
| Donor | AVI | 209 /<br>465 | 0.74 (0.14–<br>2.18), N | 6.53 (4.03–<br>12.7), N | 56.7 (30.9–<br>179), N | 0 (0–0.157),<br>S | not<br>reached |
| Donor | SpliceAI | 206 /<br>465 | 1.95 (1.06–<br>3.54), N | 9.71 (5.55–<br>23.3), N | 47.0 (26.2–<br>152), N | 0.080 (0–<br>0.183), S | 0.024 (0–<br>0.054), S |
| Acceptor | AVI | 117 /<br>13,055 | 5.69 (1.21–<br>11.3), N | 37.6 (26.0–<br>53.7), S | 669 (384–<br>1,630), S | 0 (0–0.197),<br>S | not<br>reached |
| Acceptor | SpliceAI | 115 /<br>12,923 | 2.45 (0.82–<br>4.50), N | 21.4 (13.0–<br>31.1), S | 307 (215–<br>473), S | 0 (0–0.163),<br>S | 0.014 (0–<br>0.043), S |
| Near<br>intronic | AVI | 20 /<br>13,096 | 0 (0–66.2),<br>N | 105 (44.7–<br>181), S | 491 (131–<br>1,720), S | 0.166 (0–<br>0.584), N | not<br>reached |
| Near<br>intronic | SpliceAI | 20 /<br>12,978 | 0 (0–18.9),<br>N | 22.8 (5.32–<br>45.3), N | 119 (65.5–<br>190), S | 0.347 (0–<br>1.18), N | 0.127 (0–<br>0.320), N |
| Deep<br>intronic | AVI | 103 /<br>1,572 | 2.69 (0.86–<br>5.26), N | 6.10 (3.51–<br>9.77), N | 26.7 (12.1–<br>61.8), N | 0.144 (0–<br>0.355), S | not<br>reached |
| Deep<br>intronic | SpliceAI | 103 /<br>1,538 | 4.07 (1.14–<br>10.3), N | 27.4 (12.8–<br>59.3), S | 84.0 (43.8–<br>229), S | 0.123 (0–<br>0.292), S | 0.207<br>(0.110–<br>0.326), N |

### Discrimination on splicing assays

AVI ranked splice-altering variants below raw SpliceAI in five of the six main datasets (Fig. 4a; Supplementary Tables 11 and 12). In MST1R the difference was −0.007 (−0.041 to 0.026). The pooled random-effects difference was −0.089 (−0.129 to −0.049; Hartung–Knapp interval −0.159 to −0.019; I^2^ 90%). AVI did not differ from masked SpliceAI (+0.003, −0.021 to 0.027). Against CADD the pooled difference was +0.019 (−0.010 to 0.049), with large heterogeneity (I^2^ 96%): AVI had the higher AUROC on WT1, COMPASS and OpenSplice and the lower on MST1R and POU1F1. Against phyloP 100-way it was +0.040 (0.015 to 0.066). Pangolin, run on the four minigene datasets, had the higher AUROC in three of them. The pooled difference of −0.060 (−0.115 to −0.004) was not robust with four datasets (Hartung–Knapp −0.155 to 0.036). The comparison dataset MFASS agreed with the main datasets (AVI minus SpliceAI −0.091, −0.111 to −0.071).

**Figure 4.**
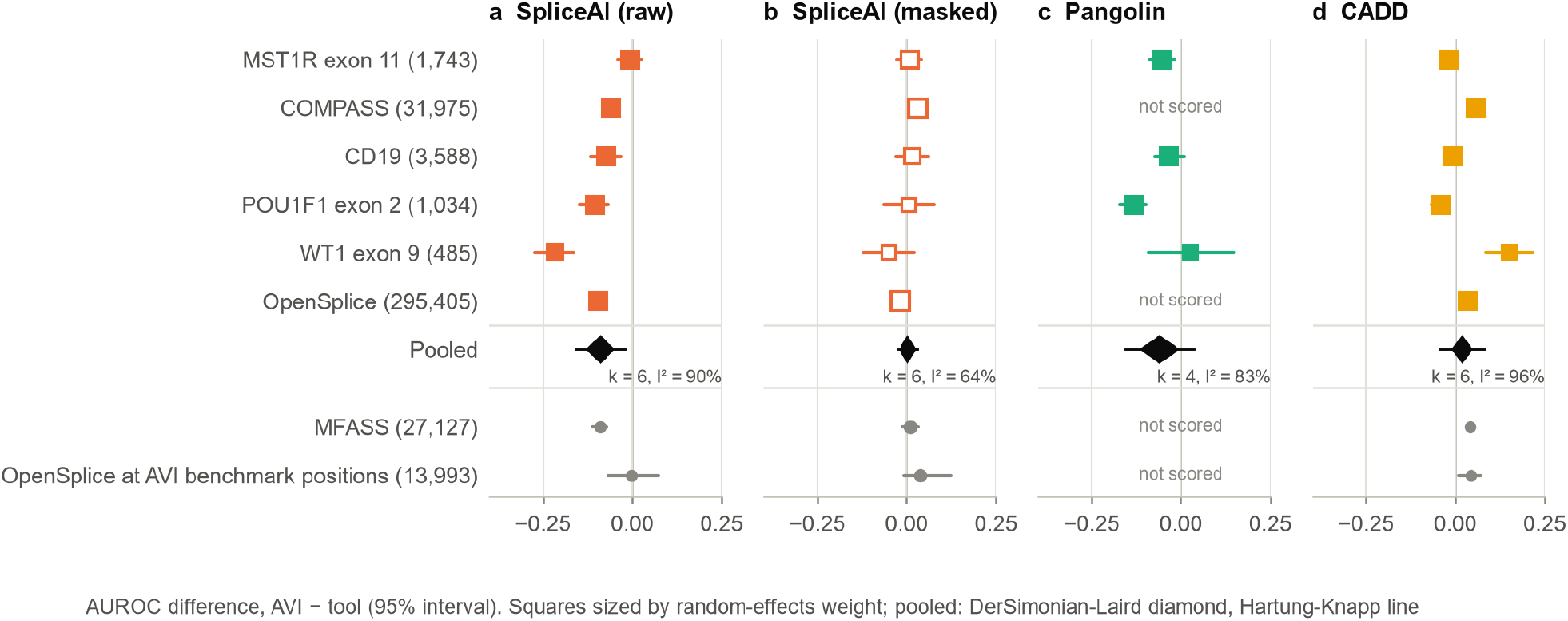
Discrimination on experimental splicing assays (Part 2). AUROC difference, AVI minus (a) SpliceAI raw, (b) SpliceAI masked, (c) Pangolin, (d) CADD, per dataset (squares, area proportional to the random-effects weight; 95% intervals from a variant bootstrap for single-gene datasets and a gene bootstrap otherwise). Number of variants in parentheses. Diamond: random-effects pooled estimate over the main datasets with its DerSimonian–Laird (DL) 95% interval; line through it: Hartung–Knapp (HK) 95% interval; k, number of datasets; I^2^, heterogeneity. Grey: comparison sets, not pooled (MFASS; OpenSplice variants in AVI developers’ benchmark data). Pangolin was run only on the four minigene datasets (POU1F1, WT1, MST1R, CD19).

AVI’s AUROC ranged from 0.522 (MST1R) to 0.837 (POU1F1). MST1R exon 11 was hard for every planned tool: AUROC 0.52 to 0.58. In OpenSplice exons in genes covered by the developers’ saturation genome editing benchmarks, analysed separately, AVI and SpliceAI did not differ (−0.001, −0.068 to 0.074).

### Additional analyses

The following analyses were added after the main results and are reported in full.

#### Top of the ranking

In the main region (Supplementary Table 13) the paired AUPRC difference between AVI and SpliceAI was +0.046 (0.002 to 0.094). The sensitivity at 99% specificity did not differ (0.777 vs 0.766; +0.011, −0.034 to 0.057), and at 95% specificity SpliceAI was higher (0.856 vs 0.899; −0.043, −0.084 to −0.003). Pangolin outperformed AVI at the top of the ranking: AUPRC −0.109 (−0.210 to −0.014) and sensitivity at 99% specificity 0.771 vs 0.893 (−0.122, −0.160 to −0.074). In deep intronic variants AVI detected 43% of P/LP variants at 95% specificity, against 80% for SpliceAI. In the predictor-not-mentioned subset the deep intronic gap at the top of the ranking remained (AUPRC −0.307, −0.399 to −0.206 against SpliceAI; −0.288, −0.483 to −0.096 against Pangolin). The AUROC gap there was smaller and its interval included zero (previous section).

#### Published thresholds

SpliceAI’s ClinGen thresholds [Walker et al. 2023] were defined on the raw score with a 4,999-base window; with the 50-base precomputed scores they held on the pooled test set (Supplementary Table 14). A score of at least 0.2 gave a likelihood ratio of 23.6 (21.9–25.4) and covered 87% of P/LP; a score of at most 0.1 gave 0.094 (0.064–0.127) and covered 91% of B/LB. The CADD thresholds recommended for missense variants [Pejaver et al. 2022] did not transfer to intronic variants (Fig. S2; Supplementary Table 14). The missense BP4 supporting interval (17.3 to 22.7) carried a likelihood ratio of 10.3 (9.09–11.7) on intronic and untranslated variants, which is pathogenic evidence above the moderate target, and it held 39% of the P/LP variants. Under the missense thresholds, an intronic P/LP variant with a CADD score of 20 would receive benign evidence. The intronic CADD thresholds derived here are 17.56, 19.18 and 23.40 for supporting, moderate and strong pathogenic evidence (Supplementary Table 6).

#### Position relative to the exon

In the assays, we classified each variant by its distance to the exon the assay measured or, in the multi-exon CD19 and MST1R minigenes, to the nearest exon of the construct (Methods; Fig. 5; Supplementary Tables 15 and 16). Pooled AVI minus raw SpliceAI was −0.161 (−0.241 to −0.080) for variants in the assayed exon (six datasets) and −0.041 (−0.062 to −0.021) for intronic variants 3 to 50 bases from the exon (six datasets). Each class pools a different set of datasets, and we did not test the classes against each other. In variants more than 50 bases from the exon, no reliable difference between AVI and SpliceAI could be shown. The DerSimonian–Laird estimate favoured AVI slightly (+0.028, 0.004 to 0.052), but it rests on two datasets, COMPASS (+0.065, −0.045 to 0.165) and OpenSplice (+0.026, 0.001 to 0.051), and the Hartung–Knapp interval included zero (−0.081 to 0.137). These variants are mostly minigene flanks within a few hundred bases of the exon, not the deep intronic variants that create pseudoexons.

**Figure 5.**
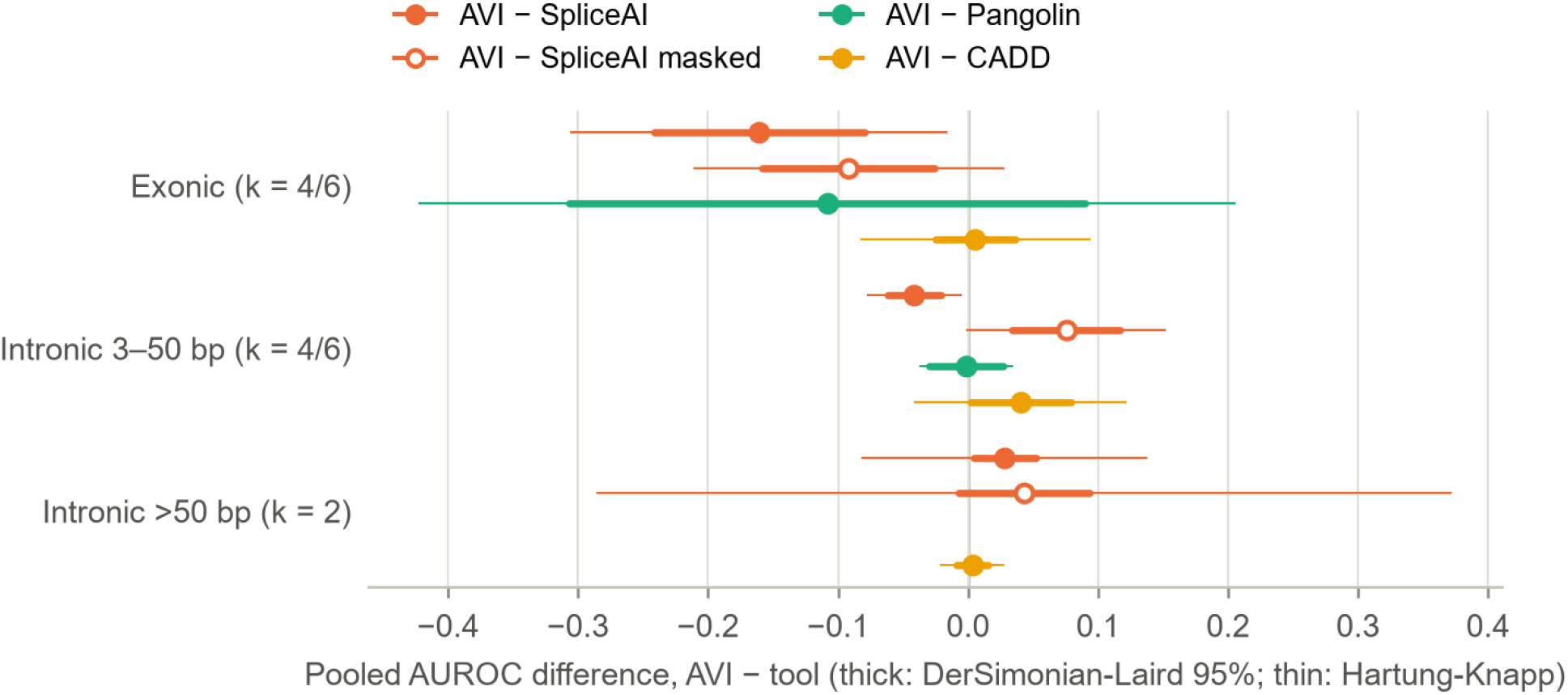
Part 2 by position relative to the exon (additional analysis). Pooled AUROC difference, AVI minus each tool, for variants in the assayed exon, intronic variants 3–50 bases from the exon, and intronic variants more than 50 bases from the exon (the Part 1 deep intronic cut). Positions come from each assay’s own description of its construct; in the two multi-exon minigenes (CD19, MST1R) intronic distance is to the nearest exon of the construct, and variants inside their other exons, beyond the outer end of a gene’s first or last exon, or within 2 bases of an exon are not shown. A dataset’s class is included when it holds at least 10 splice-altering and 10 other variants; k is the number of datasets (Pangolin / other tools). The >50-base class has two datasets (COMPASS and OpenSplice), so its Hartung–Knapp intervals are wide, and Pangolin, run only on the four minigene datasets, is absent. Thick lines: DerSimonian–Laird (DL) 95% interval; thin lines: Hartung–Knapp (HK) 95% interval.

#### SpliceAI2

SpliceAI2 [Jaganathan et al., unpublished] became public during the study (code on GitHub on 2026-10-07, version 2.0 released on 2026-10-08). We learned of it after the first Part 1 results and added it under a rule fixed before its scores were retrieved; because that decision was taken with the AVI and SpliceAI results known, it is reported here rather than among the main comparisons (Fig. S1). On new clinical labels it reached an AUROC of 0.973 (0.962–0.982) in the main region. AVI trailed it (−0.016, −0.027 to −0.006), also in deep intronic variants (−0.073, −0.120 to −0.030) and in the predictor-not-mentioned subset (−0.015, −0.028 to −0.004). Its calibration supported moderate (12.3, 6.94–22.0) and strong (219, 84.2 to unbounded) pathogenic evidence in deep intronic variants. On the assays it was ahead of AVI in every main dataset. The pooled difference over five datasets was −0.135 (−0.178 to −0.092; Hartung–Knapp −0.201 to −0.070); OpenSplice, on which its developers evaluated it, was left out of the pool and showed −0.107 (−0.123 to −0.090). In assay variants more than 50 bases from the exon, SpliceAI2 led AVI in COMPASS (−0.056, −0.108 to −0.004), the only dataset in that class once OpenSplice was set aside as developer-evaluated. SpliceAI2 had the highest AUROC in five of six main datasets; on POU1F1, Pangolin was higher (0.968 vs 0.954).

#### COMPASS without its ExAC subset

This analysis (Supplementary Table 17) was added after a review comparing the manuscript with the analysis code, and was defined before it was computed. The analysed COMPASS set holds the authors’ ExAC subset (3,746 variants, 1,213 splice-altering), chosen by presence in a population database, and 28,229 other variants (5,850 splice-altering), which approximate the authors’ random subset (Methods). Without the ExAC subset, AVI minus raw SpliceAI in COMPASS was −0.057 (−0.083 to −0.030), against −0.060 in the whole set. With COMPASS restricted in this way, the pooled difference was unchanged: −0.089 (−0.130 to −0.047; Hartung–Knapp −0.159 to −0.018) against raw SpliceAI and +0.003 (−0.021 to 0.026) against masked SpliceAI. In the ExAC subset, reported descriptively and not pooled on its own, AVI minus raw SpliceAI was −0.089 (−0.118 to −0.059). Discrimination was lower in the ExAC subset than in the other variants for every tool (AUROC: AVI 0.591 against 0.699; SpliceAI 0.685 against 0.756; masked SpliceAI 0.594 against 0.665; CADD 0.558 against 0.639; phyloP 100-way 0.529 against 0.627).

## Discussion

On ClinVar labels added after the release its developers used, AVI separated pathogenic from benign intronic variants no differently from SpliceAI and Pangolin, within about 0.02 AUROC, and better than CADD. The lead over CADD agrees in direction with the developers’ report [Cheng et al. 2026]. It held in every intronic region with enough benign variants and in every supporting set. We compared AVI with one genome-wide model only, and CADD has used SpliceAI and MMSplice scores as features since version 1.6 [Rentzsch et al. 2021]. CADD’s lag is therefore not for want of splicing input, and our comparison does not isolate what AVI’s AlphaGenome features contribute.

AVI did not outperform the splicing predictors on AUROC. It did not differ from them in the main region and trailed them in deep intronic variants. At the top of the ranking (Additional analyses) the picture was mixed: AUPRC slightly favoured AVI over SpliceAI, sensitivity at 95% specificity favoured SpliceAI, and Pangolin led AVI on both. On experimental splicing labels, AVI ranked splice-altering variants lower than raw SpliceAI and did not differ from masked SpliceAI. Because masking removes predicted changes that strengthen annotated splice sites or weaken unannotated ones, part of raw SpliceAI’s advantage on these assays may come from changes of that kind, for example variants that increase inclusion of the assayed exon; we did not test which changes AVI’s score reflects. The assay result matters because no predictor chose which variants were measured, so a gap there cannot come from the way clinical variants are found. The assay results agree in direction with the regional Part 1 results, where the splicing tools led in deep intronic and acceptor variants, though not in the main region overall.

The deep intronic gap on clinical labels needs care. Deep intronic variants usually reach a P/LP classification after a splicing predictor or an RNA study draws attention to them. Labels obtained that way favour SpliceAI-like tools, and SpliceAI and Pangolin share that advantage because they are similar splice-site models trained on similar data. Our results point both ways. The point estimate of the gap was larger where a submission mentioned a predictor (−0.108) than where none did (−0.040), which fits a contribution from how the variants were found. But the second estimate is still more than half of the full gap, its interval is wide because the subset is small, and the gap recurred in the older labels of the calibration sample (−0.043). In the assays, no reliable difference between AVI and SpliceAI could be shown more than 50 bases from the exon; those variants, however, sit in minigene flanks rather than deep in introns, come from two datasets, and SpliceAI2 led AVI on those in COMPASS (Results). The evidence is therefore consistent with part of the clinical deep intronic gap reflecting ascertainment, without showing how much. A submission that does not mention a predictor may still rest on one, and older labels may be shaped in the same way. The question needs deep intronic variants identified without predictors, for example from RNA sequencing of patient samples.

As a research measurement, AVI score intervals carried pathogenic likelihood ratios up to the strong level on the pooled test set of non-canonical intronic and untranslated variants, almost all of them intronic.

The developers call for calibration standards for non-coding variants [Cheng et al. 2026], and this is one such measurement. Three limits apply. First, AVI’s benign evidence is thin. At the ClinGen prior, AVI’s supporting benign level covered 24% of benign variants, against 90% for SpliceAI, and AVI never reached a moderate benign level, so a low AVI score carries much less information against a splicing effect than a low SpliceAI score. Second, the pooled pathogenic levels did not hold up region by region. The moderate and strong levels held in acceptor-region and near intronic variants, but no pathogenic level in donor-region or deep intronic variants, whereas SpliceAI’s moderate and strong levels held in deep intronic variants (Table 2), as did SpliceAI2’s (Results). Thresholds derived over all intronic variants should therefore not be carried into a single region without a check there, for either tool. Third, AlphaGenome and AVI outputs are provided for research use and not for diagnostic decisions [Cheng et al. 2026]; the levels reported here are measurements, not recommendations for clinical use. The CADD result shows the cost of carrying thresholds across variant classes: missense thresholds applied to intronic variants would have given benign evidence to 39% of the pathogenic variants in our set.

SpliceAI2 appeared while this study was under way and was ahead of AVI on both kinds of ground truth, including the calibrated deep intronic levels. Its evaluation is preliminary, and its terms label it for research use only, so its calibrated levels are a comparison, not a recommendation for clinical use. Its manuscript is unpublished, and its developers evaluated it on OpenSplice and on rare-disease variants from the Genomics England cohort, which may overlap with ClinVar. We report it because it shows that splicing-specific models remain ahead of a general-purpose variant score on splicing.

Several features of the design limit what the study can show. The new labels are later classifications, not independent of everything AVI saw. AVI’s frequency training can include these alleles. The exclusion of the developers’ rare-disease cases covers one submitting organisation, and the developers’ list of those variants is not public, so some overlap cannot be excluded. The calibration adapts the procedure of Pejaver et al.: it lacks their requirement that each window hold 3% of a scored reference set of rare gnomAD variants, because we had no such set. Its thresholds are specific to this calibration sample. Pangolin intervals condition on the sample of benign variants we scored and may be slightly narrow.

The statement that no predictor chose the assay variants applies to the choice of variants within the assayed exons, not to the choice of exons. OpenSplice chose exons partly by their ClinVar content. COMPASS targeted its random variants to exons and introns holding at least one ClinVar variant. Because the authors’ label for these random variants also covers their ClinVar variants, the random variants could be separated only approximately, although none of them was in the ClinVar releases of July 2024 or July 2025 (Methods). COMPASS’s ExAC subset was chosen by presence in a population database, which AVI’s frequency training may reflect, but leaving that subset out did not change the pooled result (Results). The assays measure variants in minigene constructs or libraries, whereas the tools score the genomic context.

## Conclusions

On ClinVar classifications made after the release its developers used, AVI separated pathogenic from benign intronic variants better than CADD, as its developers reported. It did not differ detectably from SpliceAI or Pangolin in intronic variants at least 3 bases from an exon, and trailed them in deep intronic variants. On splicing assays whose variants no predictor chose, it ranked splice-altering variants below raw SpliceAI. Its pathogenic evidence levels held on the pooled test set but not in every region, and its benign evidence reached 24% of benign variants, against 90% for SpliceAI. SpliceAI2, which appeared during the study and was added as a comparator after the main results, was ahead of AVI on both kinds of ground truth. How much of the deep intronic gap reflects the way those variants were found remains open until deep intronic variants identified without predictors are available. On this evidence, AVI does not replace a splicing-specific model for splicing questions, and its evidence levels, like those of any tool, should be checked in the region where they are applied.

## Methods

### Analysis plan

The question, variant sets, regions, tools, metrics and analyses were written in a study design committed to the project repository before any score was retrieved; the Part 2 plan was committed before any Part 2 score. The repository and its full commit history are public (https://github.com/aistanbulresearch/alphagenome-avi-benchmark); the analysis is at tag analysis-v1, and the two analyses added after it (COMPASS without its ExAC subset, and the ClinVar check of the COMPASS random variants; Supplementary Table 17) are at tag manuscript-v1. Variant counts per region, which contain no scores, were used to decide which analyses were main and which supporting, and to set the exclusion of older records of unresolved identity described below. The analyses in the plan are reported, with two exceptions: the comparison set of labelled variants outside the new-label set is represented by the calibration sample, and the planned secondary analyses of small insertions and deletions, which could not include AVI (AVI publishes scores for single-nucleotide variants only), were not run. Changes to code or method made after results had been produced are listed, with the numbers before and after, in the repository (CORRECTIONS.md).

### Part 1 variant sets

ClinVar [Landrum et al. 2020] classifications were read from the October 2026 release. A variant was a new label if it had a P/LP or B/LB classification in that release and none in the release of 2025-06-15, the release the AVI developers used, or in the archived releases of July 2024 and July 2025 used to trace history. Variants were excluded when they lay in benchmark data the developers used for training, selection or evaluation, and when submitted by the organisation that supplied the developers’ GREGoR cases. For TraitGym and ncVarDB, the positions of every listed variant were excluded. For the saturation genome editing screens, the positions of every scored variant were to be excluded, or the whole gene region (the gene’s GRCh38 locus with 1,000 bases on each side) where a score file could not be obtained or a scored variant had no GRCh38 position. In this analysis the gene region was used for every screen, so all variants in 13 genes were excluded (BRCA1, BRCA2, BAP1, RAD51C, VHL, DDX3X, RNU4-2, ATM, BARD1, PALB2, XRCC2, SFPQ and CTCF). Variants at the position of an older ClinVar record of at most 50 bases whose identity could not be linked to any current record were also excluded, because they may be the same variant under an older identifier. This rule was set after variant counts and before any score was retrieved; the earlier, stricter rule, which also excluded variants inside longer older records of unresolved identity (for example large structural records), is reported as a supporting analysis. Records that ClinVar had merged into newer ones were traced through its replacement history, so that a classification under a new identifier was not counted as new.

Regions were assigned from the HGVS description on the MANE Select transcript (MANE v1.5) [Morales et al. 2022]. Canonical sites are ±1 and ±2. The donor region covers +3 to +6, the acceptor region −3 to −20, and near intronic variants +7 to +50 and −21 to −50. Deep intronic variants lie beyond 50 bases. Untranslated and synonymous variants were also classified. The main region pools donor, acceptor, near intronic and deep intronic variants (intronic, at least 3 bases from the exon).

The main set comprised new labels that were single-nucleotide variants, had at least one review star, had an HGVS description on the gene’s MANE Select transcript, lay on the developers’ validation or untouched chromosomes, and were rare in gnomAD v4.1 [Guez et al. 2026] (joint exome and genome sites files, read by indexed range requests). Chromosome groups follow the developers’ data split [Cheng et al. 2026], in which training used variants on chromosomes 1, 4, 7, 8, 10, 13 and 15 and validation used variants on chromosomes 2, 5, 11, 14, 17, 20, 22 and X; the remaining autosomes (3, 6, 9, 12, 16, 18, 19 and 21) are called untouched here. A variant was rare if its joint filtering allele frequency (fafmax_faf95_max_joint) was below 0.001, the threshold used in AVI training; if that field was empty, the joint allele frequency (AF_joint) was used. Variants without a gnomAD v4.1 record were treated as rare. To check that absence was not a failed lookup, we re-queried random samples of 300 absent variants from the study set and 300 from the calibration sample (seeds 20261009 and 7): none was found in gnomAD. Two variants in each table were present in gnomAD without either frequency and were counted as rare.

Supporting analyses repeated the main comparison:

- on all frequencies;
- on rare variants present in gnomAD; on the training chromosomes;
- on variants with at least two review stars;
- with a stricter exclusion of variants inside longer unresolved older records;
- by whether any ClinVar submission of the variant mentioned a splicing predictor (a fixed list of names and phrases, such as SpliceAI, Pangolin, MaxEntScan, MMSplice or “in silico splicing”, given in the repository; presence only, no values);
- on the calibration sample.

### Scores

AVI was the raw score of the AVI scorer (AVI_SCORE) from the AlphaGenome Atlas interface (Python client 0.9.0, retrieved 2026-10-07) [Cheng et al. 2026]. At 301 positions outside the study sets, its values agreed with the downloadable AVI file (largest absolute difference 0.000366; 300 of 301 within the file’s printed precision). SpliceAI was the maximum of the four delta scores of the precomputed files for the Ensembl MANE v1.4 annotation (raw, unmasked, 50-base window; masked scores as a secondary form), taking the maximum over gene entries and over duplicate rows of an allele [Jaganathan et al. 2019]. For deep intronic variants, SpliceAI 1.3.1 was also run locally on GRCh38 with its built-in annotation and a 4,999-base window, the form for which the ClinGen thresholds were defined. Pangolin 1.0.2 [Zeng & Li 2022] was run locally with its published weights and the GENCODE v38 annotation (maximum absolute splice score change, masked, 50-base window). It was run on every non-canonical intronic and untranslated P/LP variant of the main set (461) and on a sample of 4,000 B/LB variants of the same regions, stratified by region, and inverse-sampling weights entered every metric and bootstrap. CADD was the PHRED score of the version 1.7 whole-genome file [Schubach et al. 2024]. Conservation scores were phyloP 100-way, and AVI’s own conservation inputs, phastCons 470-way and Cactus 241-way phyloP, read from the UCSC hg38 bigWig files. A variant a tool could not score was counted in a coverage table, and every comparison between two tools used the variants both scored. All scores were retrieved in one pass for all sets. The planned distance baseline was the distance to the nearest exon boundary, taken from the HGVS offset, with smaller distances ranked as more likely pathogenic.

### Discrimination

AUROC and AUPRC were computed per tool, set and region. The main comparison was the paired AUROC difference, AVI minus SpliceAI, in the main region of the main set. AVI was also compared with every other score in every region that score covered, except the distance baseline, which is reported on its own, as planned. Intervals were 95% intervals from 2,000 bootstrap replicates that resampled genes (seed 20261007), because variants of the same gene are not independent. Regional differences and intervals are not adjusted for multiple comparisons; the main comparison is the main region. Percentages are rounded to whole numbers from exact counts, except values below 1%, which are given to two significant figures.

### Calibration

Thresholds were derived with a procedure adapted from Pejaver et al. [Pejaver et al. 2022]. The calibration sample held labelled variants outside the new-label set: non-canonical intronic and untranslated variants with at least one star, every P/LP variant and 40,000 B/LB variants drawn at random, stratified by region and chromosome group, then restricted to rare variants (list committed before any score). Of its 44,001 rows, 36,191 had an earlier P/LP or B/LB classification (at or before the 2025-06-15 release or in the archived releases), 7,792 lay in the developers’ benchmark data or came from the excluded submitter, and 18 lay at older records of unresolved identity; none is in the main set, on which the thresholds were tested. AVI may score variants from its developers’ data optimistically, which could place its thresholds lower than a derivation without them; testing on the main set protects the reported likelihood ratios, and in this sample AVI and SpliceAI did not differ in the main region (AVI minus SpliceAI +0.005, −0.002 to 0.013). At up to 400 score points (quantiles of the tool’s scores), the local positive likelihood ratio was computed in the smallest symmetric window holding at least 100 labelled variants. The classes were weighted to the prior of 0.0441, so the B/LB sampling fraction does not affect it. One-sided 95% bounds came from 10,000 bootstrap samples of the labelled variants, which matches the original description of bootstrap samples of the data set. The gene-cluster bootstrap of the test phase is our addition. A pathogenic threshold is the lowest score above which the lower bound stays at or above the level’s target; a benign threshold is defined in the mirror way. Where a higher level was not reached, the score interval of the highest level reached is open-ended. The targets were those of Pejaver et al.: 2.406, 5.790, 33.53 and 1,124. We did not apply their requirement that each window hold at least 3% of a scored reference set of rare gnomAD variants, as no such set was scored here.

Each evidence level was tested within its own score interval, as evidence is assigned in the ClinGen framework; the likelihood ratios of all variants beyond each threshold, specified in our initial plan, are given in Supplementary Table 8. On the pooled test set they give the same verdicts as the interval form for AVI, SpliceAI (raw and masked), SpliceAI2, CADD and phyloP 100-way, and differ only at the supporting pathogenic level of phyloP 241-way and phastCons 470-way, whose supporting intervals are narrow or empty. Intervals were from a gene bootstrap. Where a derivation window or a test interval held no P/LP or no B/LB, the bootstrap cannot vary the empty class; we used exact Clopper–Pearson bounds of the two proportions (Bonferroni-combined) there, for every tool [Clopper & Pearson 1934]. A level was considered supported when the 95% interval of its likelihood ratio lay above the target (below the reciprocal for benign levels). Canonical and synonymous variants were not calibrated. Pangolin, scored only on a sample of the main set, entered discrimination only. Thresholds for priors of 0.01 and 0.10 used targets from the formula of Pejaver et al.

### Part 2

Assay tables were harmonised to GRCh38, and every reference allele was checked against the genome sequence; an allele listed more than once within a dataset was kept once, as its first record in the source’s own variant order (the rule does not look at labels or effect sizes; it applied to 36 OpenSplice and 108 COMPASS SNVs, of which 14 alleles in each dataset had different calls in their records; Supplementary Table 3). Labels were the authors’ own calls, or the authors’ threshold applied to their effect sizes; each dataset’s threshold and, where used, its definition of intermediate variants are listed in Supplementary Table 3 and in the repository. Intermediate variants (POU1F1, WT1) were left out of the binary metrics.

COMPASS entered, as planned, with its ExAC subset (the authors’ label) and its random variants. The authors’ label for the random variants also covers their ClinVar variants. We took those present in the ClinVar release of 2025-06-15 as the ClinVar subset and set it aside, together with the small subset the authors describe as variants from Geuvadis, a population RNA-sequencing study (their paper does not say how these variants were chosen). The remaining random variants are therefore an approximation of the authors’ random subset. COMPASS SNVs whose reference PSI was 0 or 1 in every cell line in which they were measured were not used, as the authors also exclude such sequences when they evaluate predictors; with the alleles listed more than once, this set aside 1,534 COMPASS SNVs (Supplementary Table 3). These three rules (the ClinVar and Geuvadis set-aside, the duplicate rule and the reference-PSI rule) were fixed in the code and its data notes before any Part 2 score was retrieved.

In a check that used the variant labels and no scores, none of the 28,477 random variants (28,229 after the exclusions below) appears in the archived ClinVar variant summaries of July 2024 or July 2025. In the October 2026 release 293 appear (196 of the 28,229), all first appearing after the July 2025 release; those 196 were kept in the analysis because none is classified P/LP or B/LB in that release (146 of uncertain significance, 2 with conflicting classifications, 48 without a classification), so none is a Part 1 new label. Some variants of the ExAC subset are also in ClinVar (299 of the 3,746 analysed in the July 2025 summary); this does not affect the labels, which come from the assays, and variants in the Part 1 new-label set or the developers’ benchmark data were excluded (below).

Excluded from the main analysis were:

- variants of the Part 1 new-label set;
- variants in the developers’ benchmark data, defined as in Part 1, which removed OpenSplice exons in genes with saturation genome editing data (reported separately);
- MST1R variants whose effects were median-based estimates without a test (106 single-nucleotide variants).

Scores were as in Part 1, at the genomic position of each variant. Per dataset we computed AUROC, AUPRC and the Spearman correlation of each score with the absolute measured effect. Intervals came from 2,000 bootstrap replicates resampling variants for single-gene datasets and genes for COMPASS, OpenSplice and MFASS. Paired AUROC differences were pooled over the main datasets with a DerSimonian–Laird random-effects model [DerSimonian & Laird 1986], with the bootstrap standard deviation as the standard error.

### Additional analyses

We added the following analyses after the main results:

- paired AUPRC differences and sensitivity at 95% and 99% specificity;
- the published SpliceAI thresholds and the missense CADD thresholds evaluated on intronic variants; Hartung–Knapp intervals [Hartung & Knapp 2001; IntHout et al. 2014] and leave-one-dataset-out estimates for the pooled Part 2 differences;
- Part 2 results by position relative to the assayed exon;
- SpliceAI2 as a comparator;
- COMPASS without its ExAC subset (added after a review comparing this manuscript with the analysis code, and defined in the repository before it was computed).

For the position analysis, each variant’s distance to the exon the assay measured was taken from that dataset’s own construct description. In the CD19 and MST1R minigenes, variants inside the other exons of the construct were set apart, and intronic distance was measured to the nearest exon of the construct, as in Part 1. Variants beyond the outer end of a gene’s first or last exon were not counted as intronic. A dataset’s class entered the pool when it held at least 10 splice-altering and 10 other variants.

SpliceAI2 [Jaganathan et al., unpublished], whose code became public on 2026-10-07 and whose version 2.0 was released on 2026-10-08, was added under a rule fixed before its scores were retrieved. The score was the developers’ summary score (precomputed v2.0, retrieved 2026-10-08 from the developers’ data repository), taking the maximum over gene and strand entries. All sets were scored in one retrieval. OpenSplice was reported separately because its developers evaluated SpliceAI2 on it.

For the COMPASS analysis, the analysed COMPASS variants were split into the ExAC subset and the other variants. Each part was analysed as COMPASS was, with the gene bootstrap. The pooled AVI minus SpliceAI differences (raw and masked) were recomputed with COMPASS replaced by its variants outside the ExAC subset; the ExAC subset was reported per dataset only and not pooled on its own; in the planned pooled estimate its variants are part of COMPASS.

## Acknowledgements

We thank the authors of the splicing assays analysed here (POU1F1, WT1, MST1R, CD19, COMPASS, OpenSplice and MFASS) for making their data openly available, the ClinVar submitters and the NCBI ClinVar team, and the gnomAD consortium. We thank the developers of AlphaGenome, SpliceAI, SpliceAI2, Pangolin and CADD for making their scores or software available for research use.

## Data and code availability

Code, variant lists (including the main analysis set as a list of identities and labels), result tables and figures are available at https://github.com/aistanbulresearch/alphagenome-avi-benchmark (tag analysis-v1 for the analysis; tag manuscript-v1 for the files used for this manuscript, including the main-set list, the score source records and the reproduction commands). ClinVar, gnomAD, the assay data, and the SpliceAI, CADD and conservation scores are public. AVI scores are available from the developers: the static download under a permissive licence, the Atlas interface for non-commercial use. SpliceAI2 scores are licensed by Illumina for non-commercial research. No per-variant tool scores are redistributed; the download list records the source, date and checksum of each file.

## Supplementary figures

**Figure S1.**
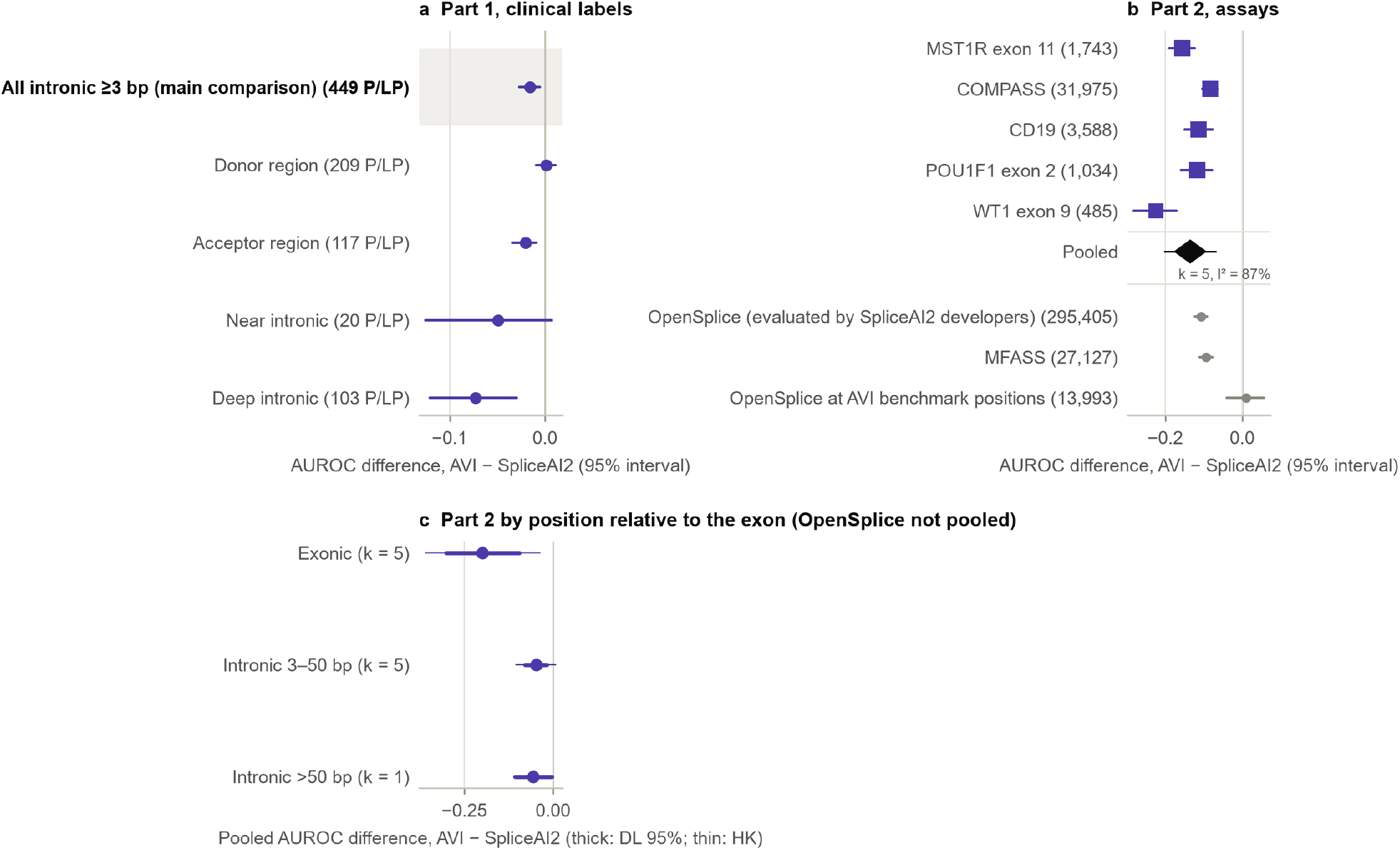
SpliceAI2 (comparator added after the main results). (a) Part 1, AUROC difference AVI minus SpliceAI2 by region. (b) Part 2 per dataset and pooled over the five main datasets; OpenSplice, on which the SpliceAI2 developers evaluated their model, is shown separately and not pooled. (c) Part 2 by position relative to the exon (OpenSplice not pooled). DL, DerSimonian–Laird; HK, Hartung–Knapp. Score: the SpliceAI2 summary score (precomputed v2.0).

**Figure S2.**
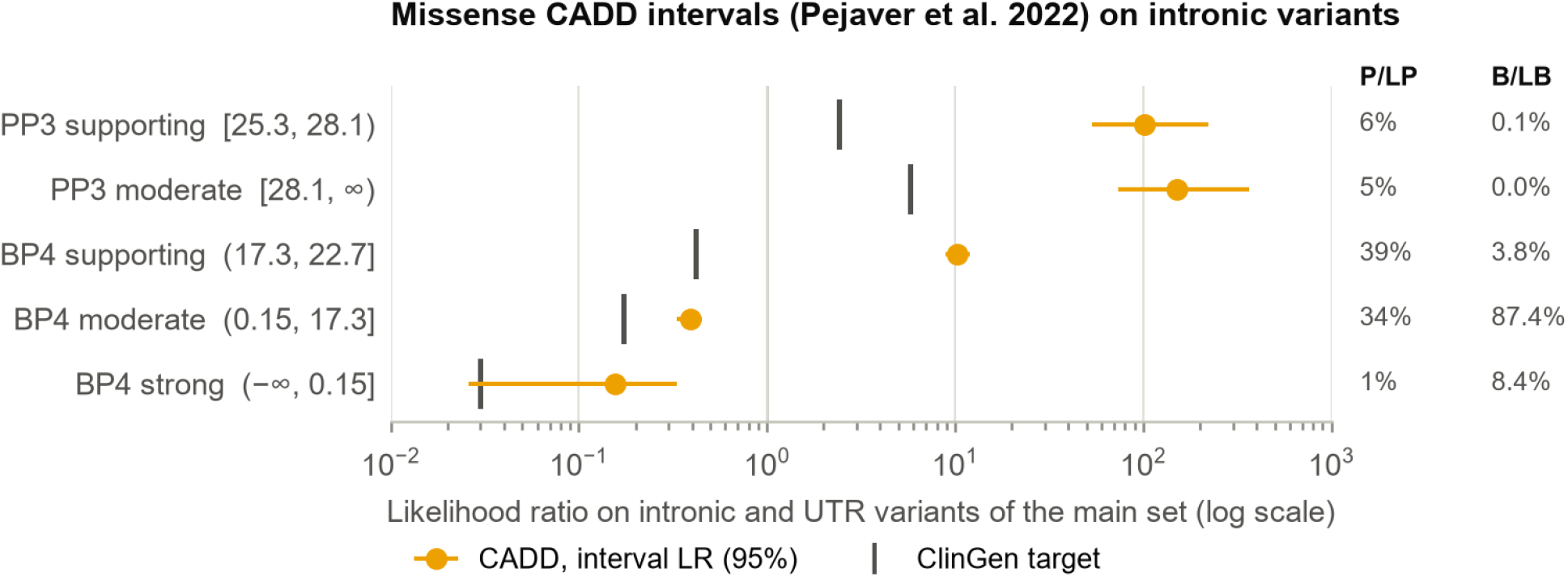
Missense CADD thresholds applied to intronic variants (additional analysis). The CADD score intervals recommended for missense variants (Pejaver et al. 2022, Table 2) evaluated as likelihood ratios on the pooled test set (the main set’s non-canonical intronic and untranslated variants); bars are the ClinGen targets for each interval’s evidence level; right, the share of P/LP and of B/LB falling in each interval. The likelihood ratios shown are in Supplementary Table 14; the intronic thresholds derived in this study are given in Supplementary Table 6.

## Notes

https://github.com/aistanbulresearch/alphagenome-avi-benchmark/tree/manuscript-v1

https://github.com/aistanbulresearch/alphagenome-avi-benchmark

### Competing Interest Statement

The authors have declared no competing interest.

